# Recurrent loss of *abaA*, a master regulator of asexual development in filamentous fungi, correlates with changes in genomic and morphological traits

**DOI:** 10.1101/829465

**Authors:** Matthew E. Mead, Alexander T. Borowsky, Bastian Joehnk, Jacob L. Steenwyk, Xing-Xing Shen, Anita Sil, Antonis Rokas

## Abstract

Gene regulatory networks (GRNs) drive developmental and cellular differentiation, and variation in their architectures gives rise to morphological diversity. Pioneering studies in *Aspergillus* fungi, coupled with subsequent work in other filamentous fungi, have shown that the GRN governed by the BrlA, AbaA, and WetA proteins controls the development of the asexual fruiting body or conidiophore. A specific aspect of conidiophore development is the production of phialides, conidiophore structures that are under the developmental control of AbaA and function to repetitively generate spores. Fungal genome sequencing has revealed that some filamentous fungi lack *abaA*, and also produce asexual structures that lack phialides, raising the hypothesis that *abaA* loss is functionally linked to diversity in asexual fruiting body morphology. To examine this hypothesis, we carried out an extensive search for the *abaA* gene across 241 genomes of species from the fungal subphylum Pezizomycotina. We found that *abaA* was independently lost in four lineages of Eurotiomycetes, including from all sequenced species within the order Onygenales, and that all four lineages that have lost *abaA* also lack the ability to form phialides. Genetic restoration of *abaA* from *Aspergillus nidulans* into *Histoplasma capsulatum*, a pathogenic species from the order Onygenales that lacks an endogenous copy of *abaA*, did not alter *Histoplasma* conidiation morphology but resulted in a marked increase in spore viability. We also discovered that species lacking *abaA* contain fewer AbaA binding motifs in the regulatory regions of orthologs of some AbaA target genes, suggesting that the asexual fruiting body GRN of organisms that have lost *abaA* has been rewired. Our results provide an illustration of how repeated losses of a key regulatory transcription factor and concomitant changes in non-coding regulatory regions of the genome have contributed to the diversity of an iconic fungal morphological trait.

**Author Summary:** Fungi exhibit tremendous variation in their asexual fruiting bodies. For example, whereas some fungi form complex fruiting bodies whose tips repeatedly generate and release spores, others produce single spores in the absence of a specialized structure. To gain insights into the molecular differences that underpin fungal asexual fruiting body diversity, we examined the genomes of hundreds of filamentous fungi for the presence of *abaA*, a master regulatory gene previously shown to control the development of fungal asexual fruiting bodies. We found that *abaA* was repeatedly lost during fungal evolution, including in a lineage of human pathogenic fungi, and that the loss of the gene was always associated with the loss of specialized structures in fungal asexual fruiting bodies. Reintroduction of *abaA* into the human pathogenic fungus *Histoplasma capsulatum*, which normally lacks the regulator, did not result in a change in the spore-producing structure but did increase spore outgrowth. Based on these results, we hypothesize that the loss of the master regulatory gene *abaA* has contributed to the observed diversity of fungal fruiting body morphology. This work advances our understanding of how fungal developmental networks evolve over time and advances our understanding of how infectious spores form in pathogenic fungi.

## Introduction

Gene regulatory networks (GRNs) that control developmental events play a major role in how species evolve [1–4]. Small aberrations in GRNs can have significant downstream consequences for development and morphology and provide a mechanism for major phenotypic changes [1]. Although the evolution of GRNs that govern morphological diversity has been extensively studied in animals [5–7], plants [8], and budding yeasts [9,10], we still know little about how the evolution of developmental GRNs is associated with the wide range of morphological diversity in the fungal kingdom.

One of the best characterized developmental cascades in filamentous fungi is the GRN controlling asexual development in *Aspergillus* (order Eurotiales, class Eurotiomycetes, phylum Ascomycota) species [11]. Asexual reproduction in the genetic model *Aspergillus nidulans* begins with the growth of a stalk, an aerial hypha that extends from the mycelium. The tip of the stalk then begins to swell, giving rise to a specialized cell called the vesicle. The vesicle then begins to produce, through budding, cells known as metulae. At the final step, the metulae produce phialides, bubble-like structures that give rise to asexual spores in a repetitive fashion (Figure 1A) [12].

**Figure 1.**
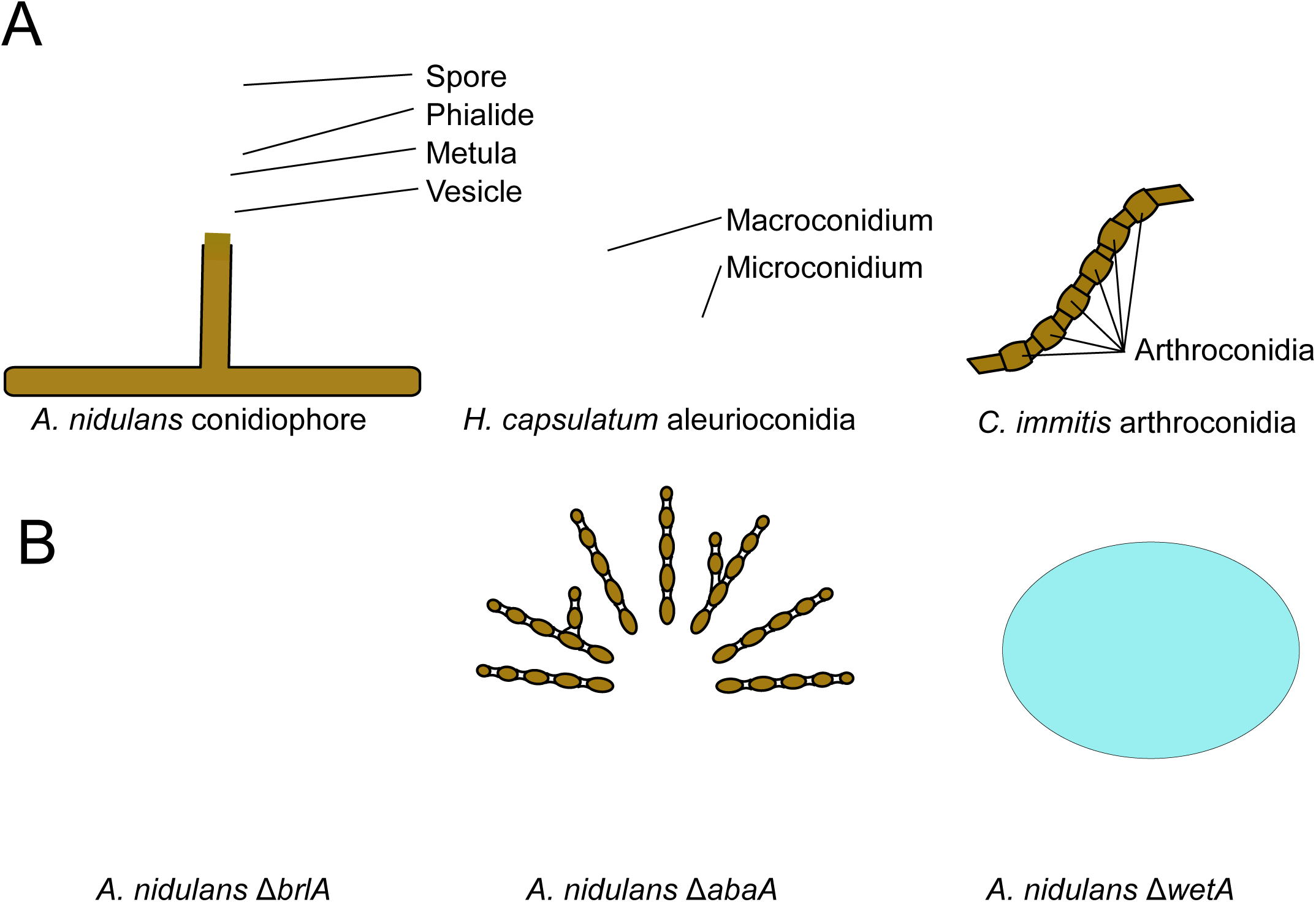
Conidial morphology differs between *A. nidulans, H. capsulatum*, and *C. immitis*, three species in the class Eurotiomycetes, and in mutants of master regulators. **(A)** *A nidulans* produces a conidiophore where spores are repeatedly generated from a phialide, whereas *H. capsulatum* produces single aleurioconidia in the form of macro- or micro-conidia and *C. immitis* forms barrel-shaped arthrocondia. (B) *A. nidulans* mutants Δ*brlA*, Δ*abaA*, and Δ*wetA* exhibit different morphological and phenotypic characteristics in asexual reproductive structures relative to the wild-type. Δ*brlA* mutants (left) form bristle-like stalks but do not form a vesicle or chains of conidia. Δ*abaA* mutants (center) are deficient in phialide differentiation, and chains of spores akin to arthroconidia form at the vesicle. Δ*wetA* mutants (right) form morphologically normal conidia, however mature conidia do not have their characteristic green color and the mutant colonies have a wet appearance.

This developmental cascade is conserved in *Aspergillus* species and their relatives, whereas the asexual fruiting bodies (conidiophores) of other organisms in the order Eurotiales or in the class Eurotiomycetes exhibit a wider range of diversity. For example, *Monascus ruber*, which also belongs to the Eurotiales, produces terminal aleurioconidia and intercalary arthroconidia. Aleurioconidia are spores produced singly from the tips of hyphae, while arthroconidia resemble barrels and develop by swelling and segmenting in the middle of hyphae. The production of both types of spores does not involve phialides or a conidiophore *per se*; instead, the mature spores form and are directly released from vegetative mycelia. Interestingly, several dimorphic human fungal pathogens in the order Onygenales, such as *Histoplasma capsulatum* and *Coccidioides immitis*, also produce aleurioconidia and arthroconidia (Figure 1A). Since infection of humans is initiated by the inhalation of spores, understanding the regulatory circuits that govern spore production in these pathogens is of significant interest.

Three DNA-binding transcription factors, BrlA, AbaA, and WetA, regulate the developmental program of the asexual fruiting body of *A. nidulans*. BrlA activates the program, AbaA regulates the development of the phialides, and WetA governs asexual spore maturation [13]. Loss of function *brlA* mutants produce “bristle-like” stalks that fail to generate asexual development structures. In Δ*abaA* mutants, the developmental program halts following the formation of metulae and mutant phialides morph into bulbous structures dispersed across rod-like metulae, giving the appearance of an abacus. Spore-producing structures appear normal in Δ*wetA* mutants; however, the resulting spores accumulate water on their surface, assume a characteristic wet appearance, and are colorless instead of the normal green (Figure 1B) [14].

The BrlA->AbaA->WetA GRN is well conserved across *Aspergillus* and related genera, as demonstrated by genetic experiments in *A. fumigatus* [15] and *Penicillium chrysogenum* [16]. Interestingly, *abaA* has been reported to be absent from the *M. ruber* genome as well as from many, but not all, genomes of species in the Onygenales [17]. The partial resemblance between the asexual structures of *M. ruber* and Onygenales species and the structures observed during asexual reproduction of Δ*abaA* mutant *A. nidulans* strains (Figure 1B), coupled with the previous reports of absence of *abaA* in *M. ruber* and some Onygenales, raise the hypothesis that loss of *abaA* is functionally linked to the simplified morphologies of the asexual fruiting bodies of these fungi.

To better understand the level of conservation of the *abaA* gene and its potential contribution to the observed diversity in asexual fruiting body morphology across filamentous fungi, we examined the distribution of *abaA* homologs across representative sets of fungal genomes. We found a direct association between the presence of the *abaA* gene and the ability of organisms to form phialides in Eurotiomycetes. Specifically, we found that *abaA* was lost in four lineages; in two of the lineages the asexual bodies do not form phialides, and in the remaining two, each represented by a single species, no asexual spores are known to form. Additionally, we observed that the Onygenales, the lineage that contains dimorphic human fungal pathogens whose members all lack phialides and *abaA*, show a decrease in the number of AbaA DNA binding motifs in the upstream regions of multiple genes orthologous to those directly targeted by AbaA in *A. nidulans*. To test if the reintroduction of *abaA* was sufficient to drive phialide formation in an Onygenales species lacking the key regulator, we ectopically expressed the *A. nidulans* copy of *abaA* in *H. capsulatum*. We did not observe changes in developmental morphology but found that total germination rates were higher in asexual spores that contained *abaA*. Our results show how GRN rewiring – i.e., repeated losses of a key regulatory transcription factor and concomitant changes in non-coding regulatory regions of the genome – has contributed to the morphological diversity of asexual fruiting bodies of filamentous fungi.

## Results

### Multiple, independent losses of the *abaA* gene in the evolutionary history of Pezizomycotina

Examination of the taxonomic distribution of the *abaA* gene across 84 species of Pezizomycotina and 3 outgroups (Table S1) identified 12 species that lacked AbaA. For the remaining 75 taxa, we identified a single copy of AbaA in at least one (75/89; 84.2%), two (72/89; 80.8%) or all three (69/89; 77.5%) of our searches. Our searches identified the *S. cerevisiae* protein Tec1 as the homolog to AbaA, which together with AbaA defines the TEA/ATTS family [18], providing additional confidence in our results. Additional searches using more relaxed cut-offs and closely related orthologous sequences as queries confirmed the presence of *abaA* in all cases. The species whose genomes lacked the *abaA* gene were *Lasallia pustulata* (class Lecanoromycetes, Ascomycota), *Bipolaris maydis* (class Dothideomycetes, Ascomycota), all nine species in the order Onygenales (class Eurotiomycetes, Ascomycota), and the outgroup species *Cryptococcus neoformans* (class Tremellomycetes, phylum Basidiomycota).

### *abaA* has been lost multiple times in Eurotiomycetes

To confirm our finding that the *abaA* gene is lost in all currently available genomes from species in the order Onygenales (class Eurotiomycetes), we conducted a more thorough search in all 154 genomes in the class Eurotiomycetes available in NCBI (as of September 17, 2017) and 10 outgroup genomes (Table S2) and mapped the presence or absence of *abaA* on the class phylogeny (Figure 2). We found that the genomes of all 31 Onygenales species examined lack the *abaA* gene. This finding is in contrast to a previous publication, which reported the presence of *abaA* in some Onygenales species [17]. Unfortunately, the previous study did not describe in detail the methods used to infer the presence of *abaA* in Onygenales, making reconciliation of their results with ours challenging.

**Figure 2.**
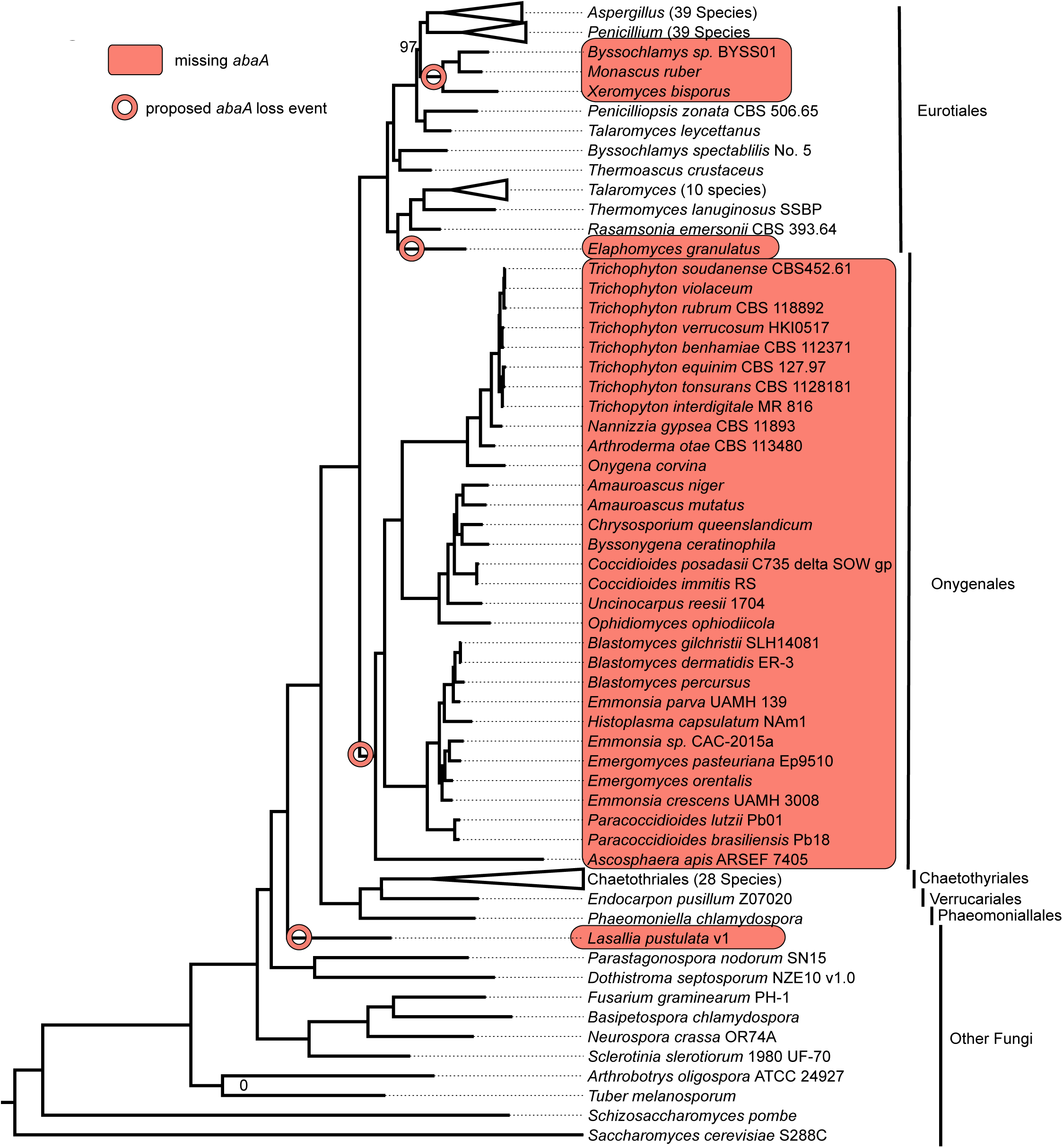
*abaA* was independently lost four times in Eurotiomycetes, including in the entire order Onygenales. Genome-scale species phylogeny of the class Eurotiomycetes with lineages missing AbaA and proposed loss events labeled. Species in the monophyletic group Monascaceae (*Byssochlamys sp. BYSS01*, *Monascus ruber*, and *Xeromyces bisporus*) and the single species *Elaphomyces granulatus* and *Lasallia pustulata* have each lost *abaA*. *Aspergillus*, *Penicillium*, *Talaromyces*, and Chaetothyriales monophyletic groups have been collapsed for ease of viewing. The full tree is available as Supplemental Figure 2. Branch labels indicate bootstrap support. Unlabeled branches have 100% bootstrap support.

The expanded set of genomes from species in Eurotiomycetes also allowed us to identify three additional *abaA* loss events; one in a common ancestor to *Byssochlamys* sp. BYSS01, *Monascus ruber*, and *Xeromyces bisporus* (order Eurotiales; Figure 2), as well as two independent losses in *Elaphomyces granulatus* (order Eurotiales) and the outgroup *Lasallia pustulata* (class Lecanoromycetes). Ancestral state reconstruction analysis of the presence/absence of *abaA* on the Eurotiomycetes phylogeny confirmed and provided strong statistical support for the inferred placements of these loss events (Supplemental Figure 3).

### Sequences similar to the AbaA DNA binding motif are less prevalent in the upstream regions of AbaA targets in Onygenales species

The inferred loss of the *abaA* gene in the ancestor of the order Onygenales led us to further investigate how the GRN controlling asexual development has evolved in Eurotiomycetes. To study possible effects of the loss of *abaA* from these genomes, we examined the distribution of AbaA binding sites upstream of *brlA*, *wetA*, *rodA*, *velB*, and *vosA* in a group of 29 Eurotiomycetes species that included 6 Onygenales species (Table S3). All five genes are known direct targets of AbaA in *A. nidulans* [19,20].

DNA binding motif searches identified 616 AbaA binding motifs in the 1.5 kb upstream region of 138 orthologs of *brlA*, *wetA*, *rodA*, *velB*, and *vosA* in Eurotiomycetes (Supplemental Figure 4). Regions upstream of *wetA* orthologs had the most motifs (average of 5.89 +/− 2.43 per region), and regions upstream of *vosA* orthologs had the fewest motifs (average of 2.44 +/− 1.34 per region). A comparison of the number of motifs in species that lack *abaA* to the number in those that possess *abaA* revealed that *brlA*, *wetA*, and *rodA* orthologs showed significantly more binding motifs in species with a copy of the *abaA* gene than in species lacking *abaA* (Mann-Whitney U tests, p-values = 1.6e-2, 2.9e-4, and 3.7e-4, respectively), while *velB* orthologs showed no significant difference (Mann-Whitney U tests, p-value = 9.1e-1), and *vosA* showed significantly more motifs in species without the *abaA* gene (Mann-Whitney U tests, p-value = 3.7e-2) (Figure 3). For example, the *wetA* ortholog present in *H. capsulatum*, a species that lacks *abaA*, has no AbaA binding motifs in its upstream region, whereas the *wetA* ortholog present in *A. nidulans*, which possesses *abaA*, has six motifs in its upstream region (Supplementary Figure 4B). These results suggest that a loss in the *abaA* gene resulted in a concomitant reduction in the number of AbaA binding sites upstream of several, but not all, of the protein’s key direct targets.

**Figure 3.**
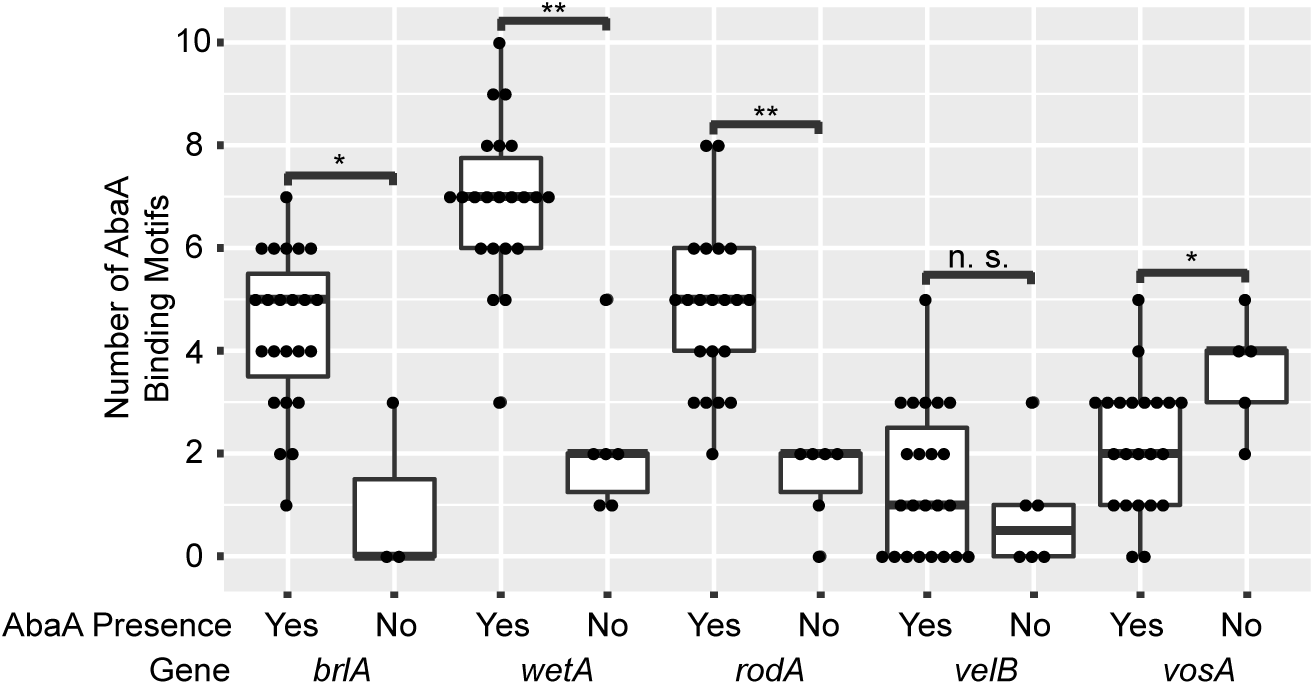
Species that lack *abaA* have fewer AbaA binding sites upstream of the protein-coding regions of many orthologous genes that are directly regulated by AbaA. Box plot comparing the number of AbaA binding sites in the 1.5 kb upstream regions of the AbaA target orthologs for species with the *abaA* gene and species without the *abaA* gene (only Onygenales species are shown here). *, p<0.05, **, p<0.01, n.s., not significant.

### AbaA is not sufficient to form phialides in *Histoplasma capsulatum*, but does increase conidial outgrowth

*Histoplasma capsulatum*, a member of the order Onygenales, lacks *abaA* and does not form phialides during asexual development. To test whether reintroduction of *abaA* is sufficient to promote phialide formation or other developmental changes, we expressed *abaA* from the model organism *A. nidulans* (*abaA(An)*) in *H. capsulatum*. We engineered the *abaA(An)* cDNA to be expressed in *H. capsulatum* under the control of the native *Aspergillus* promoter. Expression of *abaA(An)* did not lead to any obvious changes in the growth behavior of *H. capsulatum* compared to a control strain. Both strains grew with similar appearance in either yeast promoting (37°C) or conidia-producing filamentous conditions (RT) independent of the media (Figure 4A). The lack of phialides in the *abaA*-expressing mutant was further confirmed by microscopic examination of filaments that were grown on object slides covered with solid minimal medium, which induces conidia production in *Histoplasma* (Figure 4B). In addition, we examined the number of conidia produced by either the control strain or the *abaA*-expressing strain. We could not observe any significant differences between the strains, including the ratios of micro-to macroconidia in the control and the ectopic expression strains (Figure 4B, 4C). These results demonstrate that the expression of *abaA* from *A. nidulans* in *H. capsulatum* has no obvious effect on conidial morphology or levels of conidial production.

**Figure 4.**
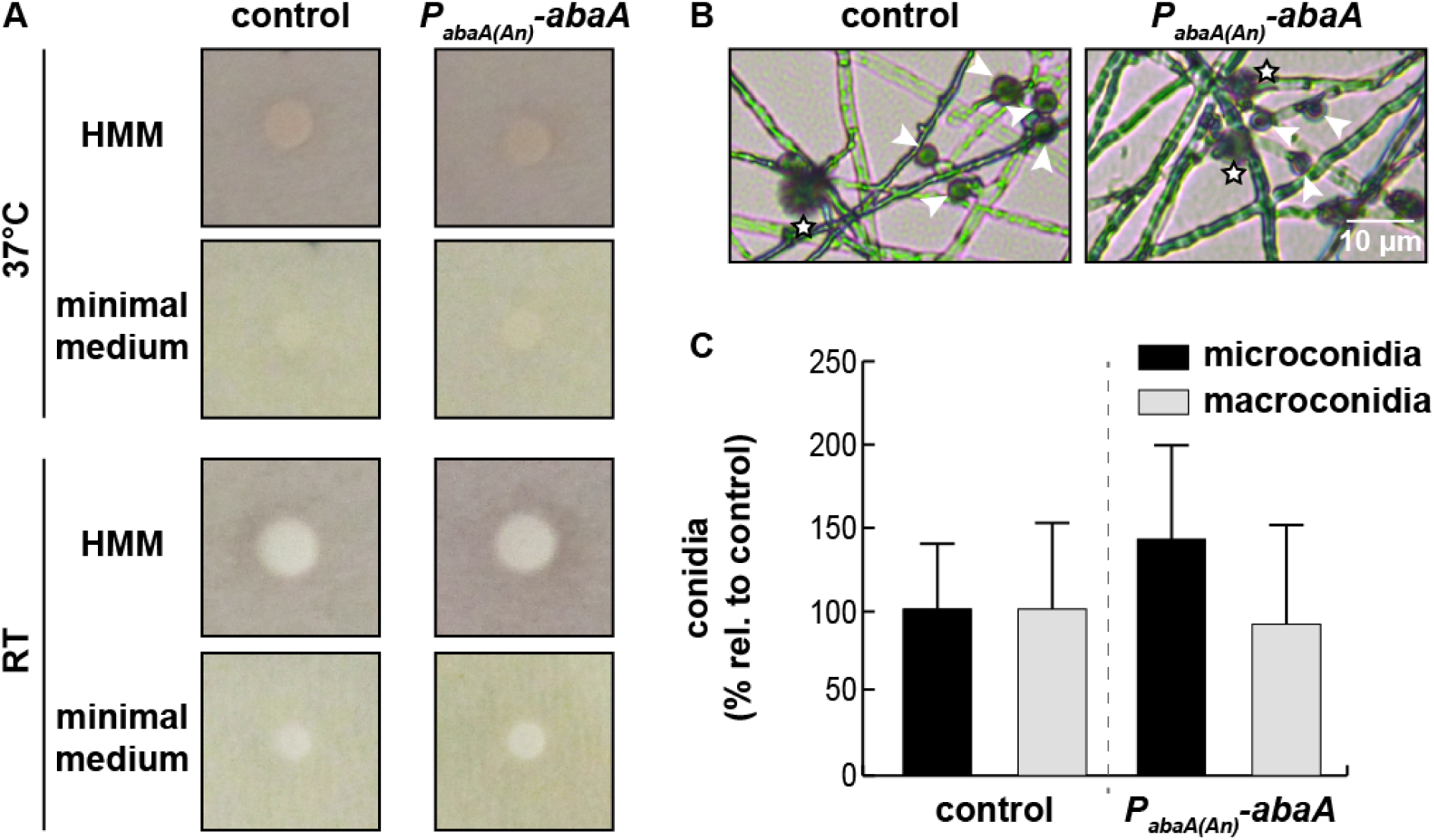
Heterologous expression of *abaA* from *A. nidulans* does not alter asexual reproductive structures in *H. capsulatum*. (A) Expression of *A. nidulans abaA* under its native *Aspergillus* promoter in *H. capsulatum* did not lead to obvious changes in growth behavior compared to a control strain, independent of the growth condition. *Histoplasma* strains were spotted on either <u>*H*</u>istoplasma <u>M</u>acrophage <u>M</u>edium (HMM) or modified minimal medium and incubated under either yeast-(37°C) or mycelial-(RT) conditions. (B) *Histoplasma* produces macro-(stars) and microconidia (white arrowheads) independent of the presence of *abaA*(*An*). (C) The total amount of conidia produced by the *abaA*(*An*) expressing strain does not differ significantly from the control strain.

Since AbaA drives the expression of genes involved in conidia maturation and viability [21–23], we tested whether the conidia produced by *abaA(An)*-containing *Histoplasma* strains differ in their ability to outgrow. Spore dilution series spotted on either HMM or minimal medium showed increased outgrowth of *abaA(An)*-derived conidia compared to the control strain independent of incubation temperature (Figure 5A). In addition, individual *abaA(An)*-derived conidia were able to form significantly more colonies on HMM at 37°C and HMM or minimal media at RT (Figure 5B). We note that no conidia were able to grow on minimal medium at 37°C, which has not been previously tested in the literature.

**Figure 5.**
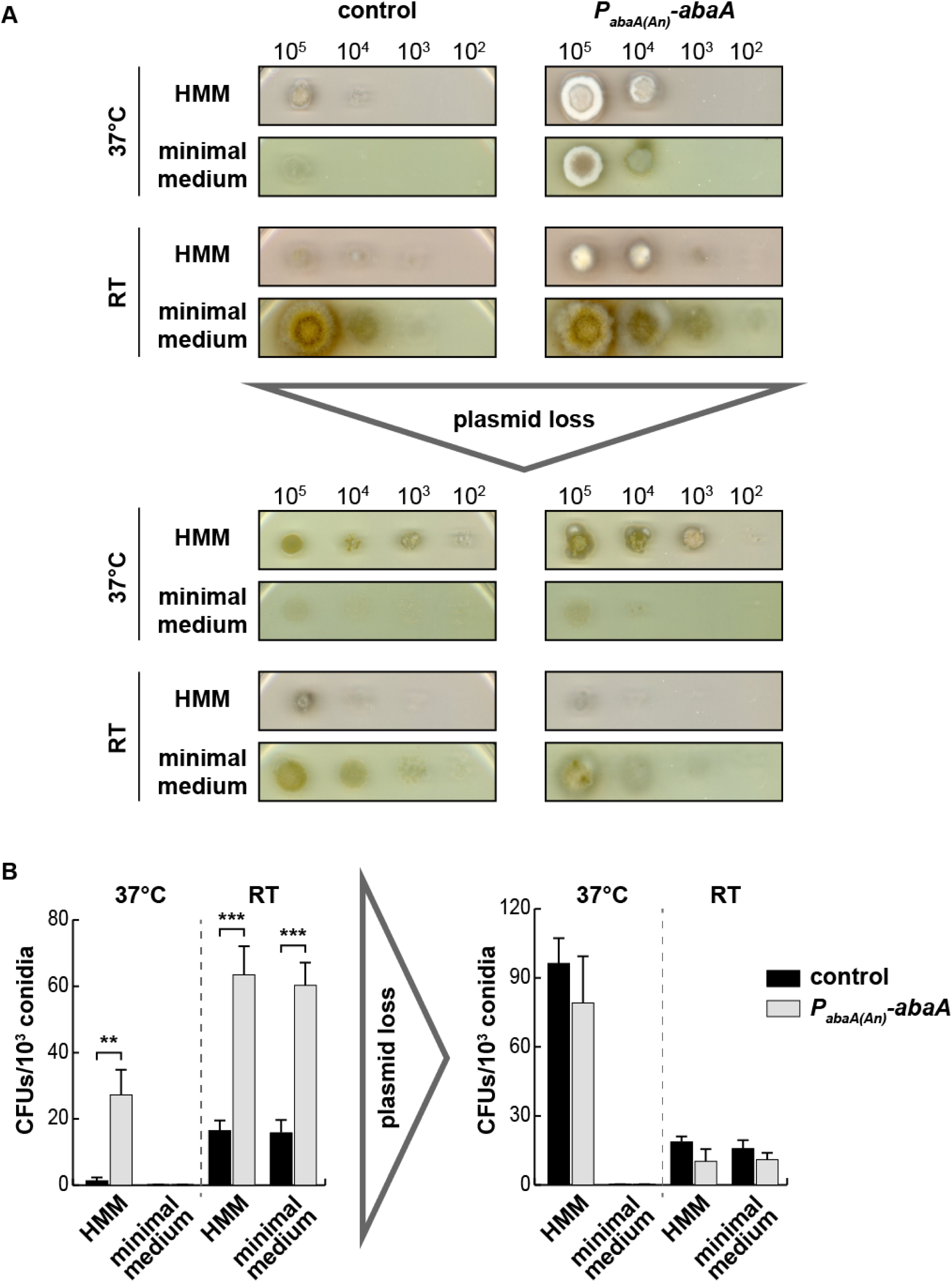
*abaA*(*An*) expression positively regulates conidia outgrowth in *H. capsulatum*. (A) Conidia harvested from *abaA*(*An*) expressing strains showed significantly better outgrowth on all tested growth conditions compared to the control. After losing both the control vector and the *abaA*(*An*) expression vector from the respective strains, outgrowth of the produced conidia was comparable between the strains, indicating that these phenotypes were dependent on the presence of the *aba*(*An*) expression construct. (B) Individual conidia derived from the *abaA*(*An*) expressing strains showed a significantly higher ability to form colonies, which was most pronounced at room temperature (RT). No significant difference was observed for CFUs of individual conidia of both strains after losing their respective vectors. Experiments were done in triplicates from two biological replicates. *P*-values were calculated using two sample *t*-test (** < 0.01; ***< 0.001).

To test whether the observed phenotypes were caused by the presence of the *abaA(An)* expression vector rather than by some genomic change that had occurred in the *abaA(An)* strains, we subjected the control and *abaA* strains to plasmid loss. Conidia derived from the isolates that had lost either plasmid showed comparable phenotypes (Figure 5), indicating that the increased fitness of conidia derived from *abaA(An)-*harboring strains was dependent on the presence of the *abaA(An)* plasmid. This result suggests that ectopic expression of *abaA* in *H. capsulatum* is sufficient to improve conidial outgrowth.

## Discussion

Transcription factors and the GRNs they control have important effects on evolution and the generation of morphological diversity. To examine the evolution of the GRN controlling asexual spore production in filamentous fungi, we determined the pattern of presence and absence for the transcription factor *abaA* across the fungal subphylum Pezizomycotina and the potential evolutionary genomic and phenotypic implications of that loss. We found that *abaA* was lost four times independently during the evolution of Pezizomycotina, including in all 31 sequenced genomes of the order Onygenales, and that this loss was correlated with differences in asexual body formation (e.g., lack of phialides in Onygenales). Characterization of the genomic effects of *abaA* loss in species that lack the transcription factor (e.g., in the Onygenales) showed that these species have experienced depletion of AbaA binding sites in the upstream regions of conserved AbaA targets. Our functional experiments showed that ectopic introduction of *Aspergillus* AbaA into the Onygenales fungus *Histoplasma capsulatum* is sufficient to impact conidial biology: conidia derived from *abaA*-containing strains showed significantly enhanced outgrowth compared to controls. Thus, it is either the case that some aspects of the *abaA*-controlled network are retained in *Histoplasma* despite the loss of the transcription factor, or that expression of *abaA* can fortuitously activate a pathway that promotes conidial outgrowth in this human pathogen.

It was notable that the expression of *Aspergillus* AbaA gave rise to enhanced conidial outgrowth in *Histoplasma* when it was expressed under the control of its endogenous, *A. nidulans* promoter. These data suggest that an unknown regulatory factor in *Histoplasma* is able to recognize the *abaA* promoter. Since *Histoplasma* has maintained BrlA, the ortholog of the upstream regulator of *abaA* in *Aspergillus*, it is possible that BrlA is driving AbaA expression and that residual aspects of the ancestral GRN are retained.

In two of the four instances of evolutionary loss of *abaA*, both of which are based on single species representatives, it is unclear if the species in question produce asexual spores at all, and if so, what form their asexual fruiting bodies take. Specifically, no asexual conidiation has been described for the lichenized fungus *Lasallia pustulata* (class Lecanoromycetes); rather, asexual reproduction in *L. pustulata* occurs by lichenized propagules which do not undergo any conidial development process [24]. Similarly, *abaA* has been independently lost in *Elaphomyces granulatus* (class Eurotiomycetes), an organism that forms symbiotic relationships and is not known to form asexual conidia [25,26]. The lack of known conidiation in both organisms is consistent with the hypothesis that loss of the *abaA* gene has an impact on conidial development.

The third instance of *abaA* loss is in the clade comprised of *M. ruber*, *X. bisporus*, and *Byssochlamys* sp. BYSS01 (Figure 2). *X. bisporus* produces “single, terminal conidia” [27], whereas *Byssochlamys* sp. BYSS01 is a species recently isolated from jet fuel for which an asexual stage has not yet been identified [28]. The final instance of loss is presumably the most ancient one, since it involves the loss of *abaA* in the entire order Onygenales (Figure 2), whose representatives are known not to form phialides during asexual reproduction [29,30]. *M. ruber* is known to form terminal conidia, without the aid of phialides [31]. A recent study showed that expressing the *A. nidulans* copy of *abaA* in *M. ruber* did not result in the production of phialides, and instead the expression of *abaA* resulted in an increase in the total number of conidia produced [32]. These results are consistent with our findings that expressing the *A. nidulans* copy of *abaA* in *H. capsulatum* results in an increase in conidial outgrowth (Figures 4 and 5).

Additionally, our findings confirm that complete loss of a transcription factor from a regulatory network is accompanied by the loss of canonical transcription factor binding sites in cognate non-coding, regulatory regions [33]. This loss of binding sites potentially reinforced the process of regulatory rewiring of the GRN following the loss of *abaA* and also helps explain why restoring *abaA* does not induce phialide formation in *H. capsulatum*.

The higher number of AbaA binding sites upstream of *vosA* orthologs in species that lack *abaA* was intriguing. A possible explanation for this gain in binding sites for a regulator that is not present is that the loss of *abaA* has forced the organisms to rewire their GRNs containing *vosA* in order to achieve compensatory functions usually controlled by *abaA*. One mechanism of this rewiring could be increasing the magnitude of *vosA* regulation by a transcription factor that recognizes sequences similar to those recognized by *abaA* by increasing the number of AbaA-like binding sites upstream of *vosA*.

The correlation between species that lack *abaA* and also do not produce phialides is striking. Therefore, we propose that the sporadic loss of *abaA* that we and others [17,32] have identified provides at least a partial, mechanistic explanation for the sporadic loss of phialide formation observed throughout Dikarya [29]. However, the selective advantage from losing *abaA* remains unclear. Both our results as well as those of Ojeda-López et al. [32] suggest that losing *abaA* (and therefore simplifying or losing the conidiophore) reduces total spore germination and production, which could be advantageous in environments where spore production may be selected against (e.g., inside a human host). Indeed, adapting to a mammalian-pathogenic lifestyle could be one case of a need for differential asexual reproduction strategies. Loss of *abaA* could also release downstream regulators from selective pressures, thus facilitating their regulatory rewiring and adoption of divergent regulatory roles. For example, in *Histoplasma*, the ortholog of WetA, the master regulator of *Aspergillus* conidial development [34,35], has been shown to regulate hyphal growth [36].

How important developmental processes and structures are controlled and evolve is a fundamental question of biology. Our examination of the repeated loss of a master regulator of asexual development in filamentous fungi and its concomitant evolutionary genomic and phenotypic changes constitute an early attempt at establishing a “fungal evo-devo” approach to answering this major question. Future studies will further reveal how changes in regulatory networks influence morphology and provide mechanistic understandings of not only the fungal lifestyle, but also the ways in which those changes dramatically impact the morphological diversity of species in the fungal kingdom and beyond.

## Materials and Methods

### Proteome and genome sequence retrieval

Whole genomes and proteomes for 84 diverse species in the subphylum Pezizomycotina (NCBI taxonomy ID 147538) as well as from 3 reference species outside Pezizomycotina (*Saccharomyces cerevisiae* strain S288C, *Candida albicans* strain WO-1, and *Cryptococcus neoformans* strain JEC21) were obtained from the NCBI FTP server on June 30, 2017 (ftp://ftp.ncbi.nih.gov/genomes/) (Table S1). Whole genomes and proteomes for the 154 species in the class Eurotiomycetes (NCBI taxonomic ID 147545) available as of September 18, 2017 as well as for 10 representative outgroups were obtained from the NCBI FTP server as well (Table S2). When multiple genomes or assemblies were available for a species, only a single representative genome or proteome was used.

### Bioinformatic searches for *abaA* in the genomes of Pezizomycotina and Eurotiomycetes

To investigate the distribution of the *abaA* gene in the genomes of filamentous fungi, we retrieved the protein sequences of *A. nidulans* AbaA and *Fusarium graminearum* AbaA from the NCBI Protein Database (GenBank accession numbers AAA33286.1 and AGQ43489.1, respectively) on August 2, 2017. blastp (from BLAST version 2.2.28) searches with each AbaA protein as a query were performed against all Pezizomycotina proteomes (Table S1) using e-value and max_target_seqs cutoffs of 1e-5 and 1, respectively. If the blastp search returned a hit, AbaA was considered to be present in that proteome.

To further confirm the presence or absence of AbaA in these proteomes using a more sensitive algorithm, we also conducted a HMMER-based search. Specifically, we obtained the seed alignment for the TEA/ATTS (TEF-1, TEC1, AbaA /AbaA, TEF-1, TEC1, Scalloped) domain (PF01285) found in AbaA proteins from the PFAM database (http://pfam.xfam.org/family/PF01285) on December 3, 2017, and used the hmmbuild tool of hmmer 3.1 (hmmer.org) to create a hidden Markov model (HMM) profile from the PFAM seed alignment. We used this profile and the HMMsearch tool of hmmer 3.1 with parameters “-E 1e-4 --tblout” to search for the TEA/ATTS domain in the 87 proteomes listed in Table S1. If hmmsearch returned any hit with these parameters, AbaA was recorded as present in that genome. To confirm losses, blastp searches were repeated on the NCBI server with the default, more relaxed parameters (e-value cutoff 10, max_target_seqs cutoffs 100) and an AbaA protein sequence from either the closest relative or a member of the nearest neighboring clade as the query. Finally, the AbaA protein from *A. nidulans* was used in tblastn (from BLAST version 2.2.28) searches performed against all Eurotiomycetes genomes (Table S2) using e-value and max_target_seqs cutoffs of 1e-5 and 1, respectively. If the blastp or tblastn searches returned a hit, *abaA* was considered present in that genome.

### Phylogenomic analysis

To reconstruct the evolutionary history of representative fungi in the class Eurotiomycetes, we employed a recently developed pipeline [37]. Briefly, we searched each proteome for 1,315 predefined universally single copy orthologous genes among 164 fungal genomes (Table S2) using the BUSCO (Benchmarking Universal Single Copy Ortholog) (version 2.0.1) pipeline [38]. The 164 proteomes contained an average of 96% +/−4% of Ascomycota BUSCO genes present in a single copy (Supplemental Figure 1A), suggesting that the selected genomes were of sufficient genome quality for phylogenomic analyses. 1,306 of the 1,315 BUSCO genes were present in at least 50% of the 164 taxa and were retained for data matrix construction (average taxon occupancy per gene: 96.31%) (Supplemental Figure 1B). Each gene was aligned using MAFFT v7.294b [39] with parameters “--reorder --bl 62 --op 1.0 --maxiterate 1000 --retree 1 --genafpair” and its gene alignment was trimmed using TrimAl 1.2rev29 [40], with the parameter “-automated1”. Alignments of all 1,306 genes were then concatenated into a single data matrix that contained 681,862 amino acid sites (average percentage of gaps: 3.77% +/− 1.09) and 164 taxa.

To reconstruct the evolutionary relationships among Eurotiomycetes, we determined the best model of amino acid substitutions using the IQ-TREE software, version 1.6.1 [41], by setting the -m parameter with the TESTONLY argument along with the parameter -mrate set to E,I,G,I+G. The best fitting model (parameter PROTGAMMAIJTTF in RAxML) was used with the RAxML version 8.2.10 [42] parameter --no-bfgs to generate 5 random trees (-d parameter) and 5 maximum parsimony trees (-y parameter). These 10 trees were used to conduct 10 independent maximum likelihood searches for the optimized tree using the parameters “-m PROTGAMMAIJTTF” and --no-bfgs; the tree with the highest log likelihood value was considered the best tree. Internode support was determined using RAxML’s rapid bootstrapping method with 100 bootstraps (parameter “-x 100”). Bootstrap values were added to the tree (RAxML parameter “-f b”) and the tree was visualized in FigTree v1.4.3 [43].

### BayesTraits analysis

To conduct analyses of ancestral state inference, we used the Bayes MultiState module in BayesTraits, version 3 [44]. We inferred the posterior probability of each character state (absence of *abaA*: 0; presence of *abaA*: 1) at the root and at each internal node of the phylogeny of Eurotiomycetes. The analysis was run for 10 million generations, sampling parameters every 4,000 generations until 2,000 samples were collected with a burn-in of the first 2 million generations. We plotted the kernel density of the posterior distribution for the posterior probabilities (PP) of states 0 and 1 at each internode and identified the largest peak values from their densities. To visualize the ancestral state at each internode across the phylogeny, we used the pie chart function in iTOL v3 [45].

### AbaA DNA Binding Motif Searches

To identify orthologs of *brlA, wetA, rodA, velB, and vosA*, known direct targets of AbaA in *A. nidulans* [19,20], in species from a subset of class Eurotiomycetes, a reciprocal best BLAST hit (RBBH) approach was used with the proteomes found in Table S3. Specifically, the proteome of *A. nidulans* was blasted against each species of interest and vice versa using an e-value cutoff of 10^−3^ and then filtered for RBBHs according to bitscore [46]. All species examined had a single significant BLAST hit for all proteins, except for *Aspergillus carbonarius* which had two significant blast hits for WetA. The two copies stem from a tandem duplication of a ~236kb region at the end of scaffold KV907500, and the sequences upstream of the two *wetA* copies were identical. The 1.5 kb genomic regions upstream of the translation start site for each of the orthologs were downloaded from either FungiDB (fungidb.org [47]) or GenBank. We used the FIMO (Finding Individual Motif Occurrences) tool from the MEME-Suite, version 4.12 [48], to locate occurrences of the AbaA binding site (5’-CATTCY-3’, where Y is either a C or a T) previously identified in the *wetA* upstream regions of *A. nidulans* [19]. Exact matches to the AbaA binding site were visualized with ggplot2 (version 2.2.1) [49].

### *H. capsulatum* strains

The generation of control and P_*abaA(An)-abaA*_ harboring strains was carried out in *Histoplasma capsulatum* G217B *ura5*^−^ background (WU15), a kind gift from William Goldman (University of North Carolina, Chapel Hill). A list of strains used in this paper can be found in Table S6. Yeast cultures of *Histoplasma* strains were propagated in liquid *Histoplasma* macrophage medium (HMM) [50] at 37°C with 5% CO_2_ on a shaker with 120 rpm. For initial phenotypic characterization of the strains, yeast cultures were diluted to an OD_600_ = 1 and 10 µl was spotted on solid HMM or minimal medium (0.5% glucose, 0.01% glutamine, 0.01% L-cysteine hydrochloride, 0.25% K_2_HPO_4_, 0.05% NH_4_Cl, 0.05% (NH_4_)_2_SO_4_, 0.01% MgSO_4_ * 7H_2_O, 0.0001% FeCl_3_, 0.1% Trace metal mix A5 with Co (Sigma-Aldrich 92949), 1% Kao and Michayluk vitamin solution (Sigma-Aldrich K3129)). These plates were incubated at either 37°C with 5% CO_2_ to remain in yeast phase mode or at room temperature (RT) in a biosafety level 3 (BSL3) facility to induce filamentous growth.

### Generation of P_*abaA(An)*_-abaA and control strains

All plasmids and primers used in this study are given in Tables S4 and S5, respectively. We created an episomal version of *abaA* cDNA under the *A. nidulans* native promoter (P_*abaA(An)*_-*abaA*) to express *abaA* in *H. capsulatum*. A *5’UTR-abaA* (AN0422) fragment was amplified from *A. nidulans* genomic DNA using primers OAS5750/OAS5749 and sub-cloned into pDONR via the Gateway cloning system (Invitrogen), resulting in pBJ212 which served as a template for further reactions. *5’UTR-abaA*_*(1. exon)*_ abaA_*(2. exon)*_ and abaA_*(3. exon)*_ were amplified from pBJ212 using primer sets OAS3322/OAS5751, OAS5759/OAS5760 and OAS5752/OAS3319 respectively. Individual fragments were fused together using primers OAS5750/OAS5749, which added attB cloning sites to the fusion product. The *5’UTR-abaA*_(cDNA)_ fragment was cloned into pDONR via the Gateway cloning system, resulting in pBJ218. pBJ218 was recombined with the episomal expression vector pSB203 that carries the *URA5* selection marker, generating pBJ226. As a *URA5* control vector we used the episomal expression vector pSB234 [51], from which we removed the Gateway cloning cassette including the *ccdB* gene for negative selection in *E. coli*, leading to pBJ238. Plasmids pBJ226 and pBJ238 were linearized to expose telomeric sequences, which facilitate episomal maintenance of the vectors in *Histoplasma*, and transformed into the G217B *ura5*^−^ parental strain via electroporation. Transformants were selected on HMM plates without uracil and verified by PCR. Strains carrying either the *5’UTR-abaA* construct or the control vector were subjected to plasmid loss by passaging the strains for three generations in non-selective HMM liquid media supplemented with uracil. Strains that had undergone plasmid loss were identified by screening for isolates that could not grow in HMM medium lacking uracil.

### Microscopy

For microscopic evaluation of control and P_*abaA(An)-abaA*_ strains, yeast cultures were diluted to OD_600_ = 0.1 and 10 µl was spotted on an object slide covered with solid minimal medium. Inoculated slides were incubated at RT for 3 weeks in a BSL3 facility. The resultant mycelium was fixed with Lactophenol blue and observed on a Leica DM1000 microscope with attached Leica DFC290 color camera.

### Conidia isolation

Conidia of *Histoplasma* strains were harvested from minimal medium plates that were inoculated with 100 µl of an OD_600_ = 1 yeast culture and incubated at RT for 4 weeks in a BSL3 facility. Plates were flooded twice with 5 ml PBS and scraped with a cell scraper to harvest conidia. The hyphal-conidia suspension was collected in a 50 ml Falcon tube. The suspension was rinsed through a syringe filled with glass wool to remove hyphal fragments and transferred to a fresh 50 ml Falcon tube. Conidia were centrifuged at 2500 rpm for 10 min, washed with 25 ml PBS, and centrifuged again at 2500 rpm for 10 min. The conidial pellet was resuspended in 500 µl PBS and stored at 4°C. Conidia concentrations were calculated using a hemocytometer.

### Conidia growth tests

We tested conidia from either control or *abaA(An)* expressing *Histoplasma* strains (and the respective strains after plasmid loss) for the ability to outgrow on different media. The indicated amounts of conidia were spotted on agar plates that were incubated at either 37°C or RT for three weeks in a BSL3 facility. To assess the capacity of individual spores to produce colonies, 10^3^ conidia of the respective strains were spread on solid media plates and incubated at the indicated temperatures for three weeks. Colony forming units (CFUs) were quantified in triplicate for each biological replicate. We note that conidia were not able to grow on minimal medium at 37°C (Figure 5B), which has not been previously tested in the literature and is of unknown significance.

## Acknowledgements

We thank Jae-Hyuk Yu (University of Wisconsin-Madison) for providing genomic DNA from *A. nidulans*. We thank Mark Voorhies and members of the Rokas lab for feedback on the manuscript and project. This work was supported by 5R01AI066224, 1R01AI146584, University of California Office of the President MRP-17-454959, and an HHMI Early Career Scientist Award (http://www.hhmi.org/research/ecs/) to AS, by a National Science Foundation grant (DEB-1442113) and a Discovery Grant from Vanderbilt University to AR, and by the Howard Hughes Medical Institute through the James H. Gilliam Fellowships for Advanced Study program (JLS and AR). AB was supported by the Vanderbilt University Summer Undergraduate Research ACCRE Scholars Program. Computational infrastructure was provided by the Advanced Computing Center for Research and Education (ACCRE) at Vanderbilt University.

## Supplementary Material

**Supplemental Figure 1. Genomes in the class Eurotiomycetes that are used in this study are largely complete.** (A) The percentage of BUSCO genes present in single copy, duplicated, fragmented, or missing in each species included in the Eurotiomycetes data set, sorted from highest percentage single copy to lowest. (B) Histogram showing the number of BUSCO genes on the Y-axis that are represented in what fraction of the genomes included in the Eurotiomycetes data set on the X-axis.

**Supplemental Figure 2. Expanded phylogeny of Eurotiomycetes.**

**Supplemental Figure 3. Ancestral state reconstruction for *abaA* loss across the phylogeny of Eurotiomycetes identifies four loss events.** Pie charts at the end of each internal branch denote the proportional value of the posterior probability (PP) of each state (Orange: absence of *abaA*; Blue: presence of *abaA*). Two types of circles next to species names are used to code trait data: circles filled with orange denote absence of *abaA* (i.e., 0); circles filled with orange denote presence of *abaA* (i.e., 1).

**Supplemental Figure 4. Positions of AbaA binding sites in the upstream regions of AbaA target orthologs.**

**Supplemental Table S1. Pezizomycotina genomes used.**

**Supplemental Table S2. Eurotiomycete genomes used.**

**Supplemental Table S3. IDs of AbaA direct target orthologs.**

**Supplemental Table S4. Plasmids used in this study.**

**Supplemental Table S5. Primers used in this study.**

**Supplemental Table S6. Strains used in this study.**

