## Supplementary figures and images for "Recurrent loss of *abaA*, a master regulator of asexual development in filamentous fungi, correlates with changes in genomic and morphological traits"

### Supplementary Figures

#### Supplemental Figure 1

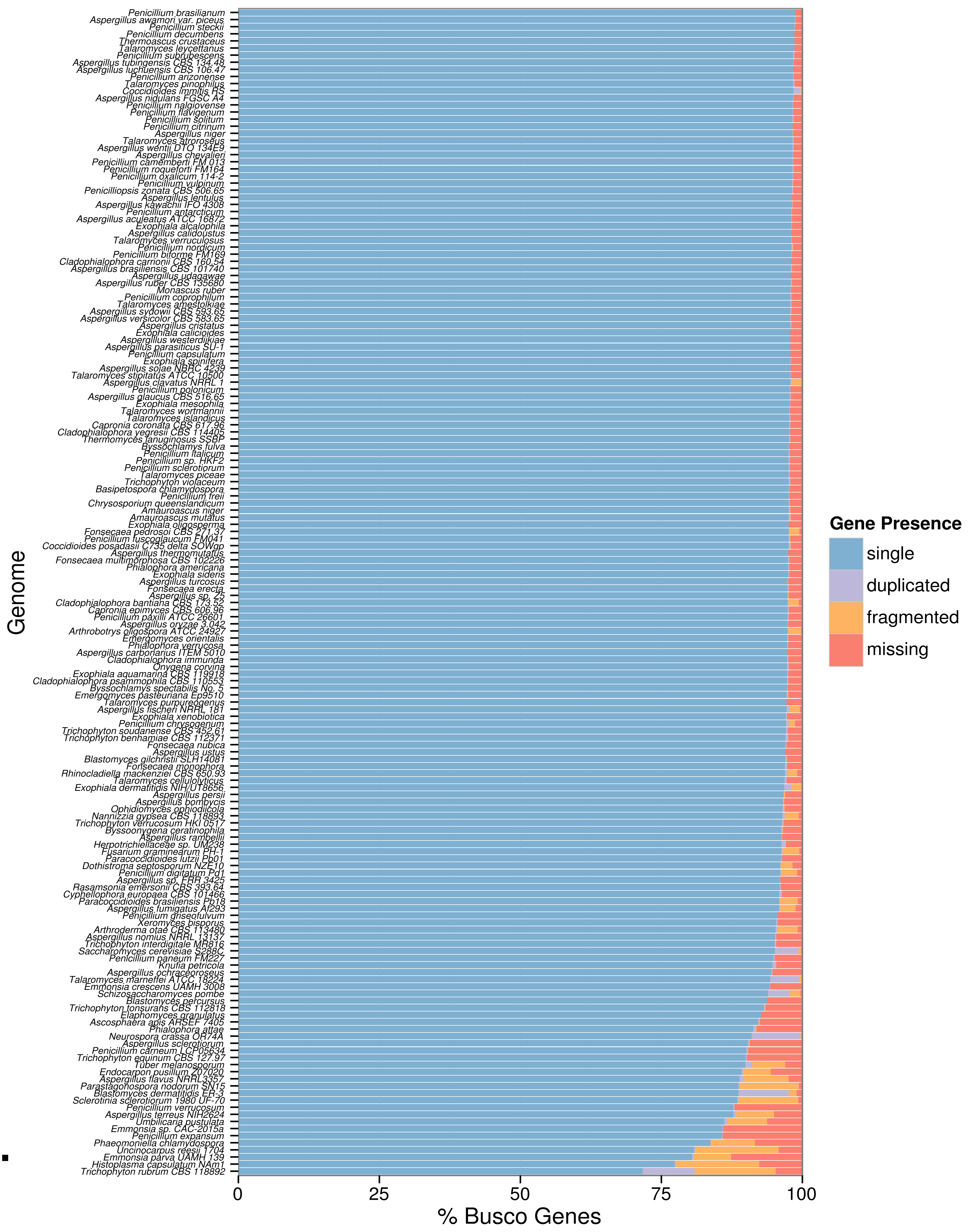


A.

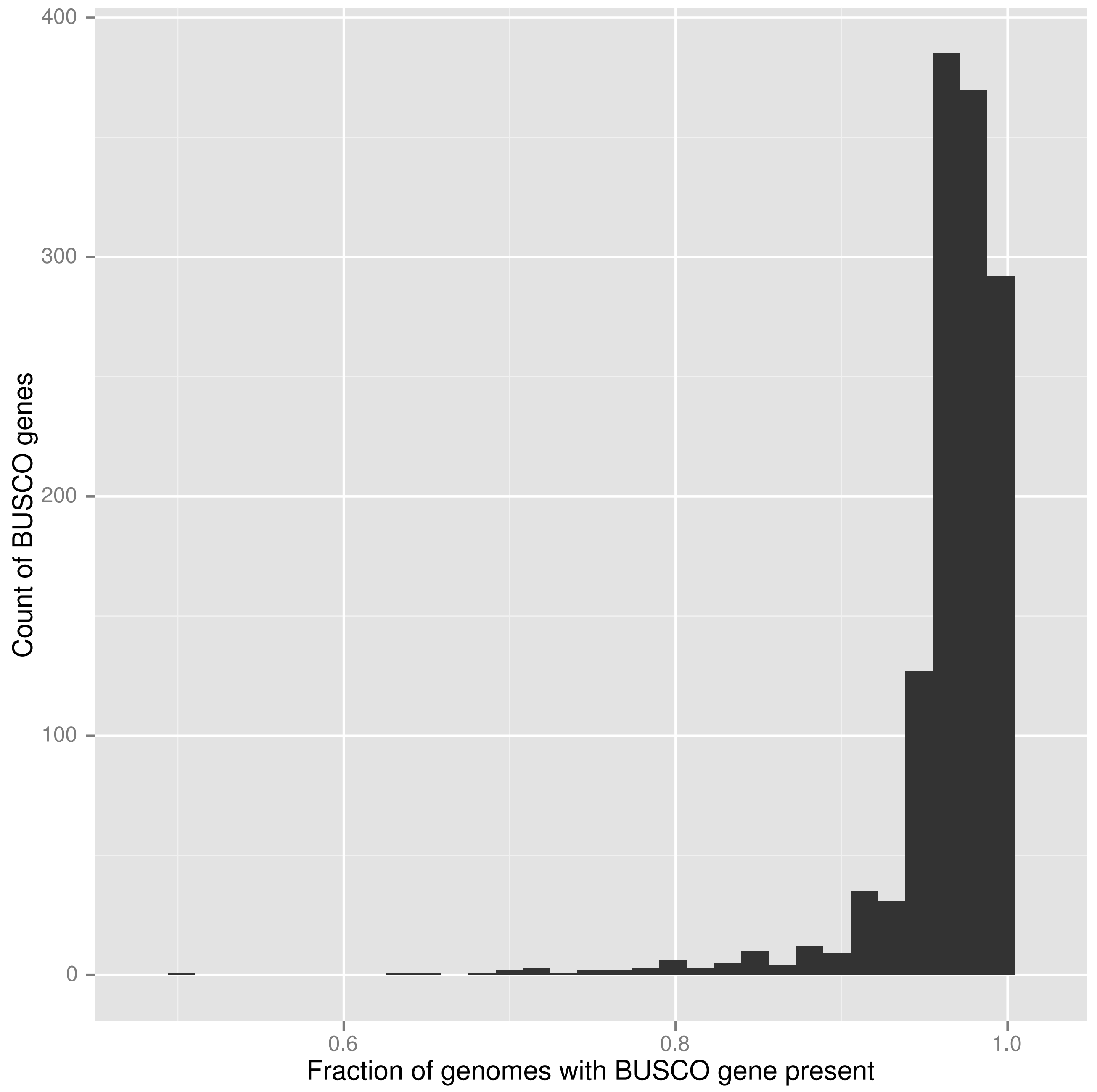


B.

Supplemental Figure 2

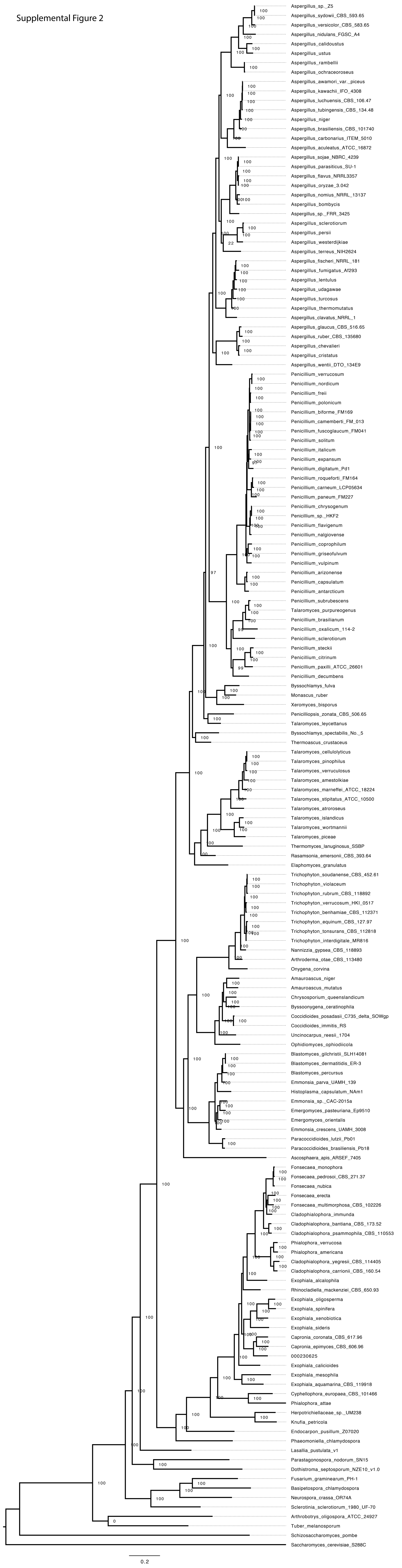

#### Supplemental Figure 3

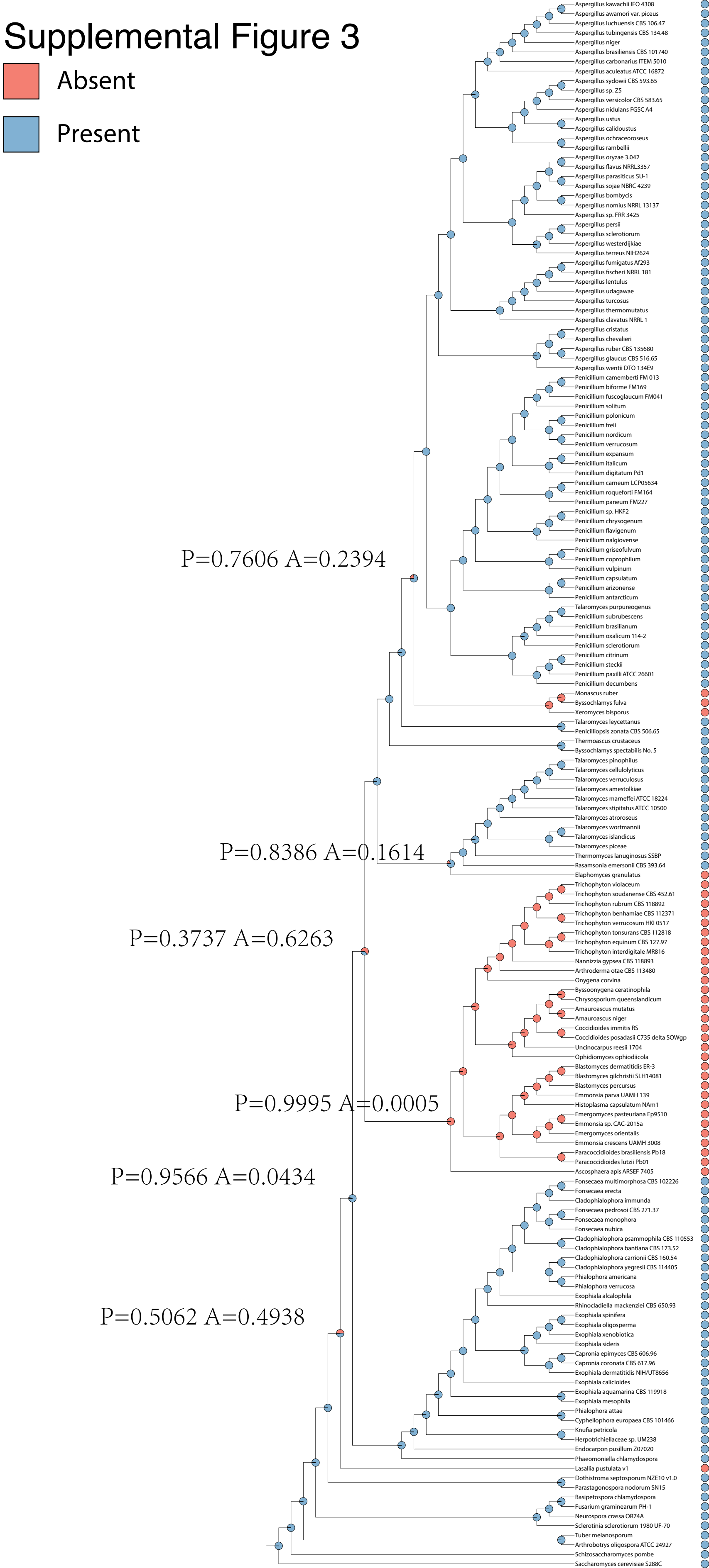

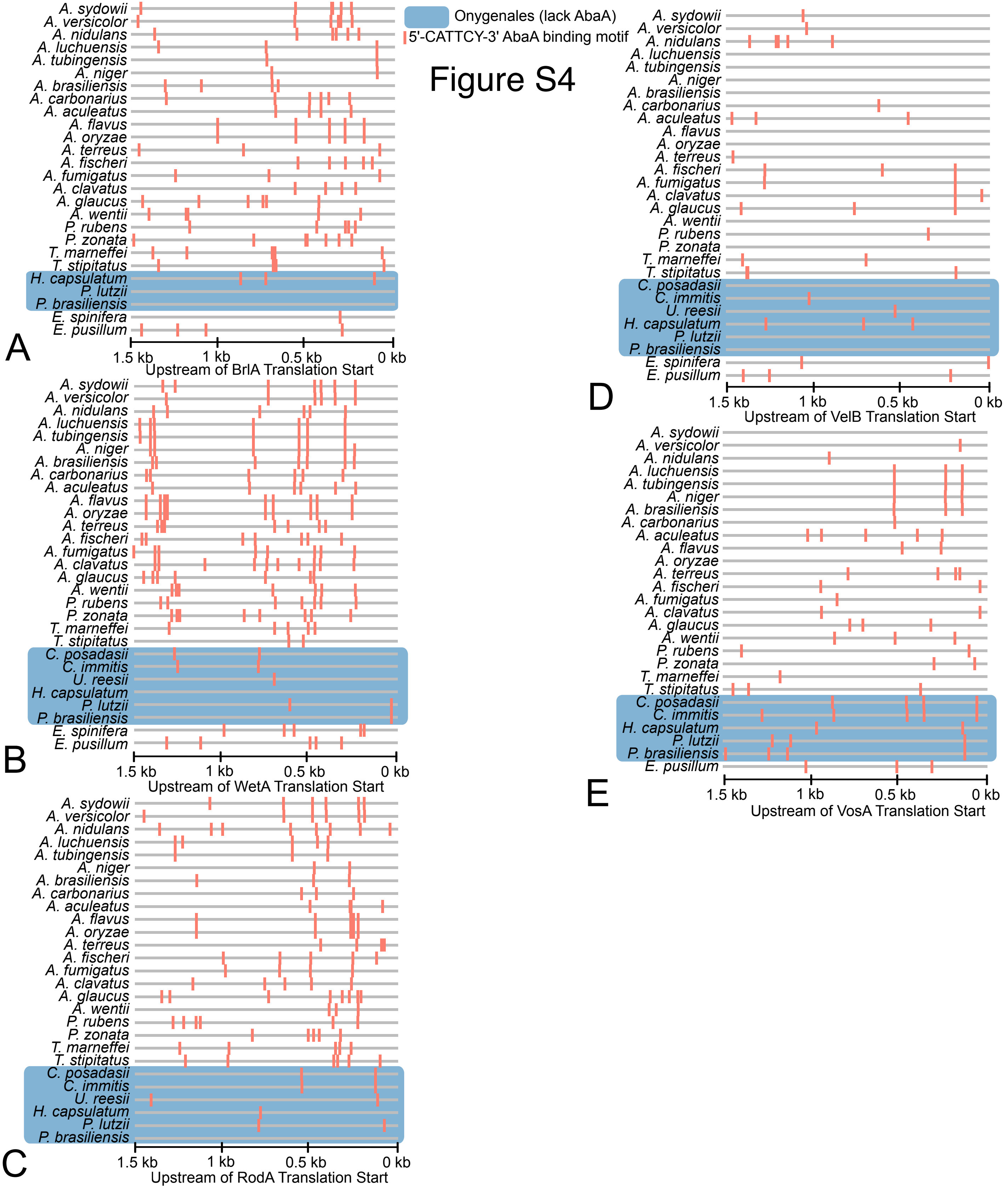
